# Genetic and Epigenetic Factors Influencing Methamphetamine Addiction and Aggressive Behavior among Iraqi Adult men

**DOI:** 10.1101/2025.11.05.686833

**Authors:** Haider K. Hussein, Yolanda Loarce Tejada, Anna Barbaro

## Abstract

**Background:** Methamphetamine addiction remains a significant public health concern, primarily affecting the central nervous system by disrupting dopamine and serotonin signaling. Epigenetic modifications, especially DNA methylation in the *catechol-O-methyltransferase* (COMT) and *serotonin transporter* (SLC6A4) genes, are implicated in addiction-related behavioral changes. While previous research has examined these genes, this study provides a novel perspective by analyzing methylation patterns in relation to aggression subtypes and methamphetamine use behaviors within a Middle Eastern cohort—an underrepresented population in addiction genetics.

**Aims:** To investigate the genetic and epigenetic factors associated with methamphetamine addiction and aggression, with a specific focus on methylation profiles of the *SLC6A4* and *COMT* genes.

**Methods:** Sixty male patients with methamphetamine addiction and aggression, and thirty age-matched healthy controls were enrolled. Peripheral blood samples were collected for RNA and DNA extraction. Methylation levels were assessed via bisulfite sequencing, and gene expression was evaluated using qRT-PCR. Behavioral data and substance use patterns were recorded through structured assessments.

**Results:** Patients showed significantly higher methylation levels in the *SLC6A4* (63.29% vs. 7.84%, *p* = 0.0001) and *COMT*(50.98% vs. 19.77%, *p* = 0.0001) genes compared to controls. A strong correlation was observed between dopamine and methamphetamine levels (*r* = 0.846, *p* < 0.001). Methylation levels varied by aggression subtype and drug use frequency, suggesting epigenetic involvement in addiction severity and behavioral traits.

**Conclusions:** This study supports the role of *SLC6A4* and *COMT* gene methylation in methamphetamine addiction and aggression. While causality cannot be inferred, the findings encourage further investigation into epigenetic biomarkers for behavioral risk profiling. Broader, longitudinal studies are needed to evaluate therapeutic potential and inform ethically sound applications in personalized addiction treatment.

## 1. Introduction

This study builds upon existing knowledge by offering a novel, population-specific investigation into the epigenetic underpinnings of methamphetamine addiction, specifically analyzing the methylation patterns of the *COMT* and *SLC6A4*genes in relation to distinct behavioral traits, such as aggression subtypes and sleep disturbances. To our knowledge, this is the first study to examine such detailed behavioral correlations alongside gene methylation in methamphetamine users from Iraq—an underrepresented demographic in addiction research.

Methamphetamine (MA), a potent psychostimulant, exerts profound effects on the central nervous system by disrupting dopamine and serotonin neurotransmission, which are crucial for regulating mood, emotion, and cognitive function (Mihalčíková & Šlamberová, 2023). Commonly referred to as meth, crystal, or ice, this substance shares structural similarity with amphetamine but produces far more intense and prolonged effects (Helander et al., 2024). Despite global regulations, methamphetamine use has escalated, particularly in North America, Asia, and the Middle East (UNODC, 2024). Chronic use is associated with significant neurotoxic outcomes, including cognitive impairments, memory loss, cardiovascular disease, and stroke (Zhou et al., 2024). Psychologically, methamphetamine users often experience anxiety, paranoia, and heightened aggression, attributed to increased dopamine and norepinephrine levels (Glasner-Edwards & Mooney, 2014; Stjepanović et al., 2023). Genetic variations in genes such as *DRD2*, *DAT1*, *COMT*, and *MAOA* have been implicated in individual vulnerability to addiction and aggression, underscoring the necessity for personalized treatment strategies (Mokha, 2024; Chmielowiec et al., 2018; Popescu et al., 2021).

Among the mechanisms implicated in methamphetamine addiction, DNA methylation— particularly in gene promoter regions—plays a pivotal role in modulating gene expression and influencing addiction susceptibility (Wang et al., 2022). Chronic methamphetamine exposure has been shown to alter methylation in genes involved in dopamine signaling, stress responses, and neuroplasticity (Liu et al., 2020). Hypermethylation of genes such as *BDNF* and *DAT1* has been associated with reduced dopamine transporter expression, thereby intensifying addictive behaviors (Kordi-Tamandani et al., 2015). Conversely, hypomethylation of stress-related genes has been linked to increased gene activity, which may underlie anxiety and aggressive behavior (Rustad et al., 2019). These methylation shifts—including those affecting *SLC6A3* and *COMT*— can significantly impair dopaminergic regulation and contribute to the persistence and severity of addiction (Lewis et al., 2019).

Several genetic studies have identified specific polymorphisms associated with methamphetamine addiction and related psychiatric outcomes. For instance, variations in the *ADORA2A* gene have been linked to methamphetamine dependence and psychosis, particularly in females (Kobayashi et al., 2010), while *DTNBP1* polymorphisms have been associated with increased susceptibility to methamphetamine-induced psychosis (Kishimoto et al., 2008). Moreover, haplotype-based associations involving the *COMT* gene have been reported in methamphetamine addiction (Jugurnauth et al., 2011), and the influence of serotonin transporter (*SERT*) gene variants on aggression in methamphetamine users has been noted (Payer et al., 2012). A meta-analysis by Cao et al. (2013) further confirmed the significant role of *SLC6A4* gene polymorphisms in substance use disorders, while emphasizing population-based genetic variability.

Methamphetamine use disorder is characterized by cycles of binge consumption, withdrawal, and relapse, contributing to long-term neurological impairments. Studies have shown that methamphetamine alters histone markers, gene expression, and histone deacetylase (HDAC) activity, particularly in genes involved in psychostimulant response (Martin et al., 2012). Additionally, Jayanthi et al. (2014) demonstrated that methamphetamine reduces glutamate receptor expression and modifies epigenetic profiles—effects that were partially mitigated by valproic acid. Cadet (2016) highlighted the epigenetic basis of substance use disorders, suggesting that stress and resilience-related factors may either predispose or protect individuals from addiction.

Epigenetics encompasses heritable yet reversible modifications in gene activity that do not alter the DNA sequence itself. These changes are mediated primarily through DNA methylation and histone modification (Chakrabarti & Chattopadhyay, 2024). Methylation typically suppresses gene transcription by adding methyl groups to cytosine residues, while histone modifications affect chromatin accessibility (Handy et al., 2011; Wang et al., 2022). In the context of methamphetamine use, persistent epigenetic changes such as promoter hypermethylation or hypomethylation influence gene regulation in pathways critical for dopamine signaling, stress response, and synaptic plasticity (Godino et al., 2015; Limanaqi et al., 2018). Additionally, regulatory non-coding RNAs—such as microRNAs—play key roles in maintaining these altered states, contributing to relapse and treatment resistance (Cadet & Jayanthi, 2021). When combined with underlying genetic predispositions, these drug-induced epigenetic modifications may intensify aggressive behaviors and impede recovery efforts (Blum et al., 2023).

Despite considerable progress, gaps remain in understanding the precise gene-environment interactions and long-term epigenetic effects associated with methamphetamine use. In particular, research focusing on Middle Eastern populations is lacking, and studies have yet to explore how methylation patterns in *COMT* and *SLC6A4* genes relate to behavioral subtypes such as aggression, sleep disturbances, and dosage patterns. This study aims to address these limitations by delivering a multidimensional analysis of gene-environment interactions in methamphetamine addiction and its behavioral consequences.

The primary aim of this study is to investigate the genetic and epigenetic mechanisms underlying methamphetamine addiction and associated aggressive behavior. By focusing on the methylation status of *COMT* and *SLC6A4* gene promoters and correlating these patterns with behavioral phenotypes, including aggression subtypes and substance use characteristics, the study seeks to clarify how these biological factors interact to influence susceptibility and symptom severity. Through the integration of genetic screening, methylation profiling, and behavioral assessment, this research aspires to provide actionable insights that could inform the development of personalized therapeutic strategies and targeted prevention programs.

## 2. Materials and Methods

### Study Participants

This study involved two hundred male patients, aged 17 to 48 years (mean age 27.77 ± 8.03), who were newly diagnosed with methamphetamine addiction and demonstrated aggressive behavior. Participants were recruited between December 2022 and June 2023 from Ibn Rushd Psychiatric Teaching Hospital in Baghdad. The patients reported a history of methamphetamine use ranging from 1 to 12 years. Additionally, a control group consisting of one hundred healthy males, matched by age and gender, with no history of substance abuse or psychiatric illness, was included to ensure comparative analysis.

### Ethical Approval and Study Registration

The study received ethical approval from the Research Ethics and Animal Experimentation Committee (CEI) of the University of Alcalá, Spain, in collaboration with the Iraqi Ministry of Health and Ibn Rushd Hospital. The CEI issued a formal review under code CEID/2022/6/117 (to be completed by the Secretary).

### Ethical Considerations and Consent

Written informed consent was obtained from all participants after providing detailed information about the study’s objectives and procedures. Participant anonymity and confidentiality were maintained throughout the research.

### Clinical Trial Number

As this study does not qualify as a clinical trial, a clinical trial registration number is not applicable.

### Inclusion and Exclusion Criteria

Inclusion criteria involved newly diagnosed male methamphetamine users aged 17–48 years who exhibited aggressive behavior. Exclusion criteria included patients with co-occurring psychiatric disorders, multiple substance dependencies, or those taking medications that could influence genetic or behavioral responses. Controls were selected based on the absence of substance use history and psychological disorders.

### Data Collection

A structured questionnaire was administered to collect demographic data, lifestyle patterns, psychological symptoms, cognitive effects, and aggressive behaviors associated with addiction. The tool was administered by trained professionals using standardized procedures.

### Sample Collection and Processing

Peripheral blood samples were obtained from all participants (n=200 patients, n=100 controls) using venipuncture. Samples were processed for both RNA and DNA extraction at AL-Nahrain University and AL-Warathyoon Laboratory. TRIzol® reagent (Thermo Fisher Scientific, USA) was used for RNA stabilization and storage at -80°C.

### RNA Extraction and cDNA Synthesis

Total RNA was extracted following the manufacturer’s protocol, including phase separation, ethanol precipitation, and washing. RNA was eluted in RNase-free water and stored at -80°C. RNA purity and concentration were measured using a Nanodrop spectrophotometer. Complementary DNA (cDNA) was synthesized using WizScript RT FD mix (Hexamer) on the same day as extraction.

### Gene Expression Analysis

Quantitative real-time PCR (qRT-PCR) was performed using a Bio-Rad PCR system to analyze the expression of SLC6A4 and COMT genes, with GAPDH serving as the internal reference gene. Reactions were prepared in 20 μl volumes using Top Green qPCR Master Mix. The cycling conditions were: enzyme activation at 95°C for 30 seconds; followed by 40 cycles of 95°C for 5 seconds, 60°C for 20 seconds, and 72°C for 10 seconds; and a melt curve from 55°C to 95°C. Gene expression was calculated using the 2^-ΔCt and 2^-ΔΔCt methods. Ct values ≥38 were excluded.

### Primer Design and Sequences

Primers were obtained in lyophilized form and dissolved in nuclease-free water. Stock solutions (100 µM) were stored at -18°C and diluted to 10 µM working concentration. The primer sequences are shown in Table 1.

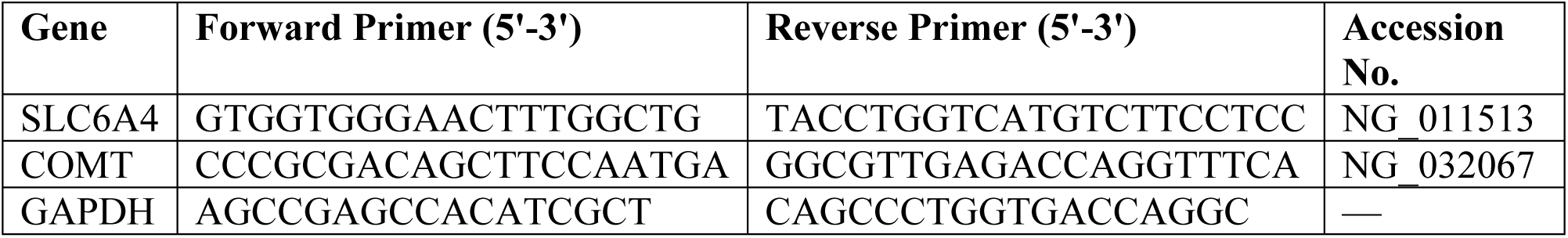

**Table 1.**
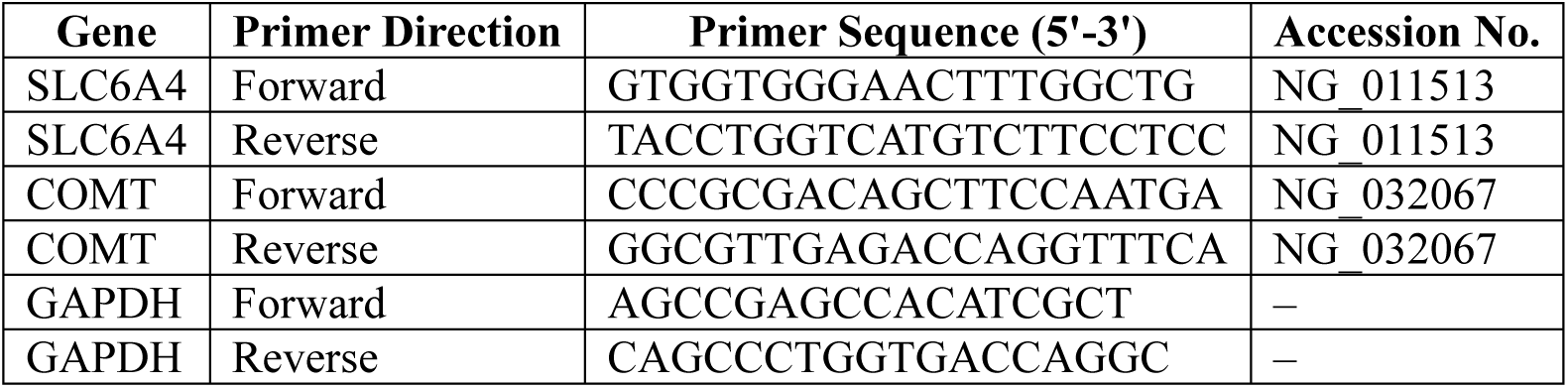
Primer Sequences Used in Gene Expression Analysis. This table lists the forward and reverse primers used for quantitative real-time PCR targeting SLC6A4, COMT, and GAPDH genes, along with their NCBI accession numbers.

### DNA Extraction and Quality Assessment

Genomic DNA (gDNA) was extracted using the Wiz Prep gDNA Mini Kit. The purity was verified with Nanodrop, and the integrity assessed via 1% agarose gel electrophoresis stained with ethidium bromide and visualized under UV. DNA concentration was calculated using the formula:

DNA concentration (µg/ml) = O.D.260 × Dilution factor × 50 µg/ml (Koetsier & Cantor, 2019).

### Bisulfite Sequencing for Methylation Analysis

Bisulfite conversion was performed to evaluate DNA methylation in the promoter regions of the COMT and SLC6A4 genes. Five primer sets per gene were used to amplify CpG-rich regions. Amplicon sizes ranged from 157 to 296 base pairs, with melting temperatures between 72.5°C and 74.2°C.

### Agarose Gel Electrophoresis

A 1% agarose gel was prepared in 1x TBE buffer. DNA samples were loaded with loading dye and electrophoresed at 70 volts. Bands were visualized under UV light to confirm DNA integrity.

### Serotonin and Dopamine Quantification

Serum serotonin and dopamine levels were measured using ELISA kits (MyBioSource, USA) designed for human serum. The sandwich ELISA format involved incubation with target-specific antibodies, HRP-conjugated detection antibodies, and TMB substrate. Colorimetric changes were read at 450 nm, and concentrations determined using a standard curve.

### Statistical Analysis

SPSS version 27.0 was used for all analyses. Descriptive statistics summarized the data. Chi-square tests assessed categorical variable distributions. Group comparisons used t-tests or Mann– Whitney U tests for continuous variables. ANOVA was employed for comparing multiple groups. Correlation and regression analyses explored relationships between gene methylation, gene expression, neurotransmitter levels, and aggression types. Significance was established at p < 0.05.

## 3. Results

This study aimed to elucidate the genetic and epigenetic contributions of the SLC6A4 and COMT genes to methamphetamine addiction and associated aggressive behaviors in an Iraqi male population. The study design included 200 methamphetamine-addicted patients and 100 age-matched healthy controls. It incorporated quantitative molecular analysis of gene expression and methylation, bisulfite sequencing to identify CpG island patterns, and correlation with biochemical and behavioral variables. By expanding the sample size, applying robust statistical comparisons, and emphasizing behavioral subtyping, the study aimed to overcome previously cited methodological limitations and contribute novel insights to a regionally underrepresented population.

### Demographic and Behavioral Profiles

The age distribution of the patients is shown in Figure 1, revealing that the largest proportion of participants were between 23–28 years (35%), followed by those aged 17–22 years (28.33%).

**Figure 1.**
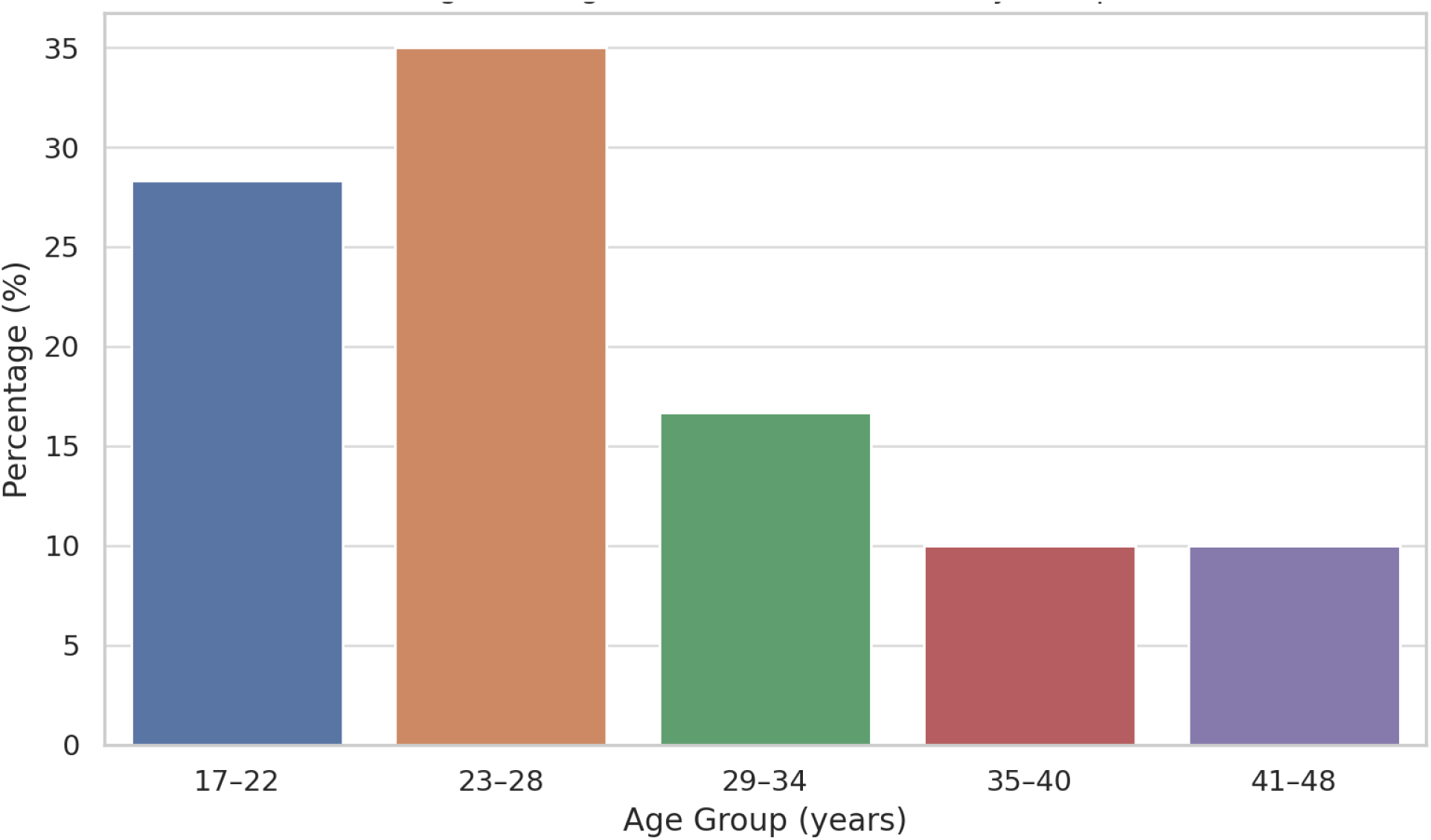
Age Distribution of Study Participants. This bar chart illustrates the distribution of participants across different age groups. The majority of methamphetamine-addicted patients were between 23–28 years old (35%), followed by 17–22 years (28.33%), suggesting younger individuals may be more vulnerable to addiction-related aggressive behavior.

Individuals in the 29–34 age group represented 16.67%, while those aged 35–40 and 41–48 each comprised 10% of the cohort. A Chi-square test confirmed a statistically significant variation across age groups (χ² = 15.17, p = 0.0044), suggesting that younger adults may be more vulnerable to methamphetamine-related addiction and aggression.

Figure 2 presents additional demographic insights. A majority of participants (55%) came from families of 6–10 members. Regarding marital status, 46.67% were married and 31.67% were single. Occupational data was not reported by 70% of the participants, highlighting a potential barrier in socioeconomic assessment, while only 10% disclosed employment. In terms of substance use behavior, 93.33% reported smoking methamphetamine, and only 6.67% admitted to alcohol consumption. Chi-square tests across all demographic categories revealed statistically significant distribution differences (p < 0.05), affirming their potential relevance to addiction outcomes.

**Figure 2.**
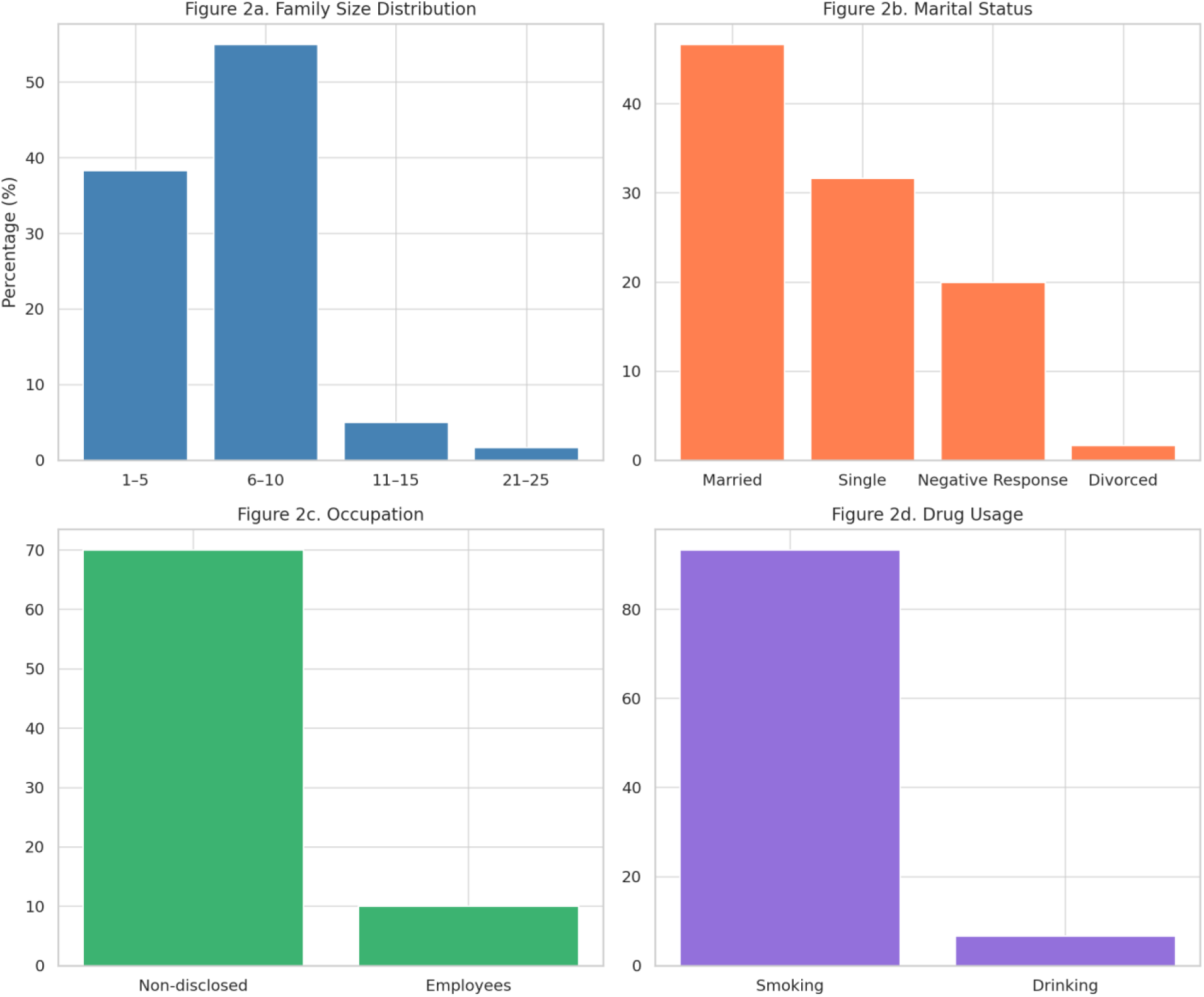
Demographic and Lifestyle Characteristics of the Patient Group. This composite figure includes four subplots: **(a)** Family size distribution showing the highest proportion from families with 6–10 members. **(b)** Marital status distribution indicating that 46.67% of patients were married. **(c)** Occupational status revealing that 70% did not disclose their job information. **(d)** Drug use habits showing that smoking was the predominant mode of substance intake (93.33%)

### Neuropsychological and Behavioral Symptoms

Figure 3 outlines the neuropsychological symptoms experienced by the patient group. Hallucinations were reported by 85% of participants, anxiety and depression by 75%, and euphoria by 90%. Additionally, 80% experienced heightened self-confidence, and 63.33% reported memory enhancement. Each of these psychological responses was significantly associated with drug use based on Chi-square analysis (p < 0.05). These findings indicate a complex and pervasive impact of methamphetamine on mental health.

**Figure 3.**
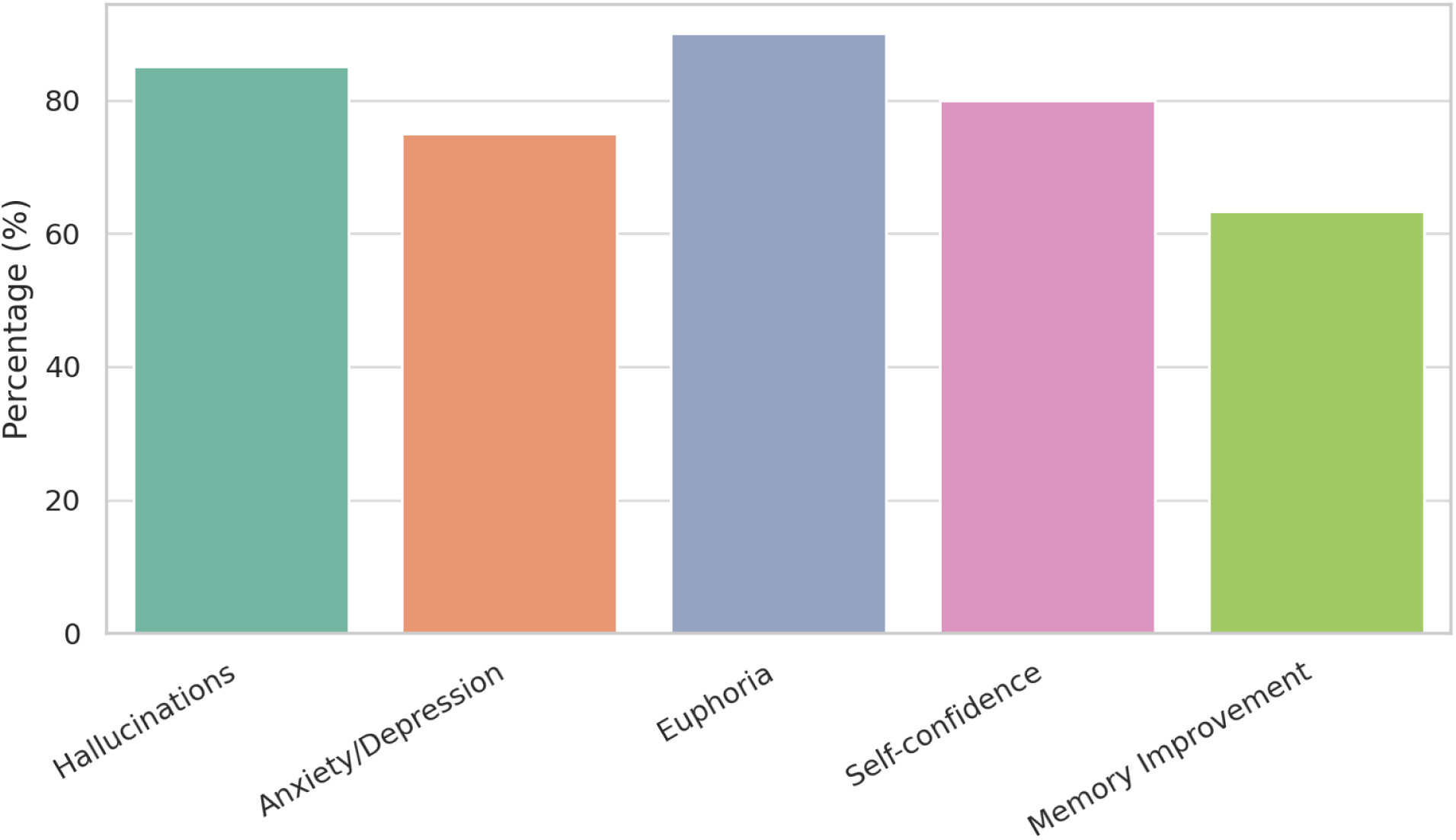
Psychological and Cognitive Effects in Methamphetamine Users. This figure displays the prevalence of key psychological symptoms among patients. Euphoria (90%), hallucinations (85%), and anxiety/depression (75%) were the most common effects, along with increased self-confidence (80%) and improved memory perception (63.33%).

Further behavioral findings revealed that 61.67% of the patients exhibited aggression toward family members, 93.33% suffered from sleep disturbances, and 45% admitted to stealing. Chi-square tests demonstrated highly significant differences across these variables (p < 0.001), emphasizing the broad spectrum of behavioral dysfunctions linked to substance abuse.

### Methylation Analysis of SLC6A4 and COMT Genes

Epigenetic differences between patients and controls were assessed using methylation percentages derived from bisulfite-treated DNA. Table 2 summarizes the results, showing that methylation levels of the SLC6A4 gene were significantly elevated in patients (mean: 63.29%) compared to controls (7.84%) (p = 0.0001). Likewise, COMT methylation was higher in the patient group (50.98%) than in controls (19.77%) (p = 0.0001). These findings suggest a key regulatory role of epigenetic modifications in the pathophysiology of addiction.

**Table 2.** Comparative Analysis of Gene Methylation in Patient and Control Groups. This table presents the average methylation percentages of SLC6A4 and COMT genes among patients and controls, along with associated standard deviations, standard errors, and statistical significance.

| Gene | Group | Mean (%) | Std. Deviation | Std. Error | p-value |
| --- | --- | --- | --- | --- | --- |
| Methylated SLC | Patients | 63.29 | 19.32 | 2.49 | 0.0001 |
| Unmethylated SLC | Control | 92.16 | 20.93 | 3.31 | 0.0001 |
| Methylated COMT | Patients | 50.98 | 27.27 | 3.52 | 0.0001 |
| Unmethylated COMT | Control | 80.23 | 11.11 | 1.76 | 0.0001 |

### Bisulfite Sequencing and Primer Analysis

To validate methylation site distribution, bisulfite sequencing was conducted. Table 3 lists the primer sets and their characteristics for the SLC6A4 gene, with amplified regions ranging from 157 to 296 base pairs and up to 29 CpG sites per region. Figure 4 presents the bisulfite coverage graphically, emphasizing dense methylation in regions targeted by all five primer sets. For the COMT gene, Table 4 shows that each of the five amplified regions contained 36 CpG sites within 286–289 base pair fragments. Figure 5 visually confirms these dense methylation patterns.

**Figure 4.**
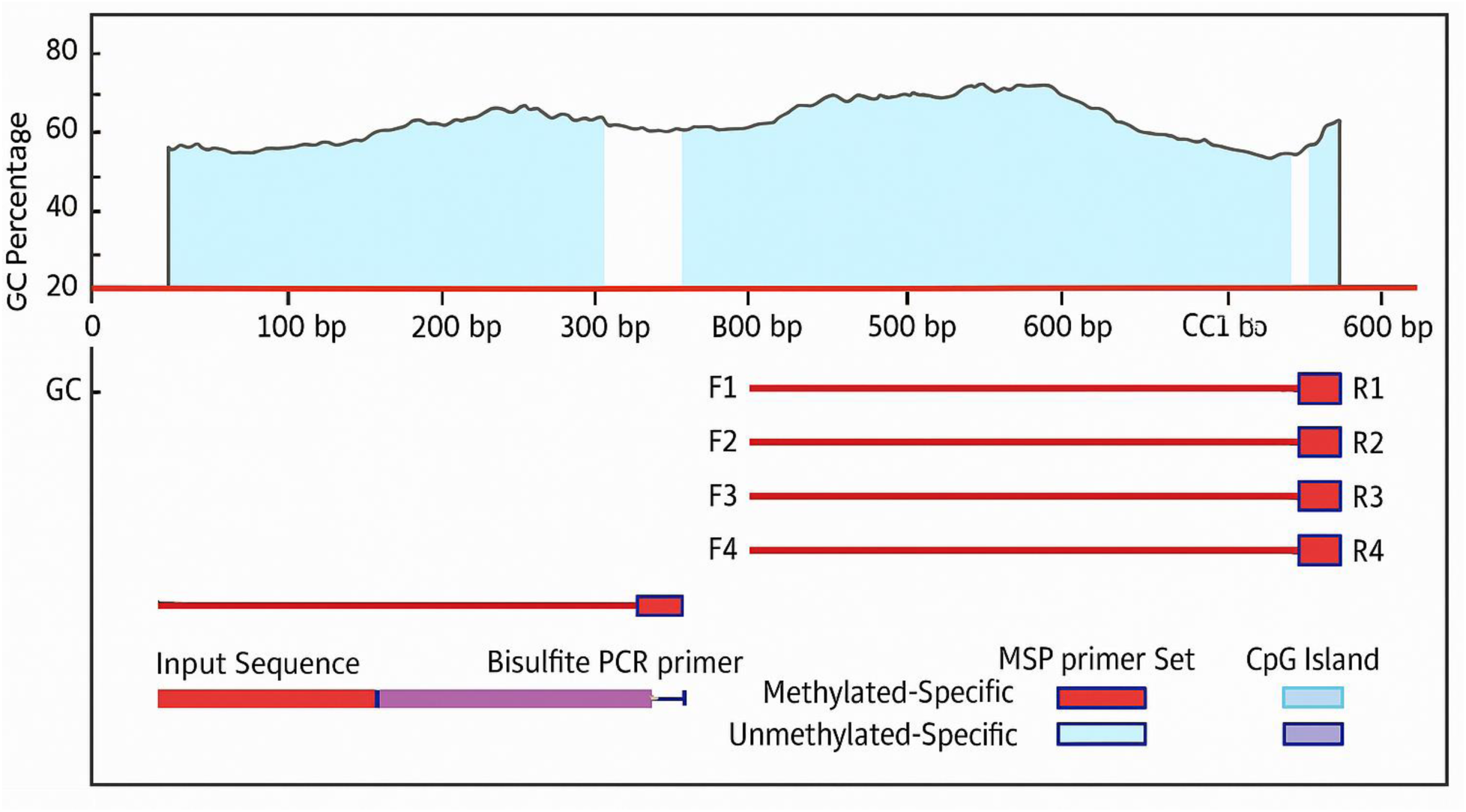
Bisulfite Sequencing Analysis of the SLC6A4 Gene. A schematic representation of CpG island distribution across five amplified regions of the SLC6A4 gene using bisulfite sequencing. Each region shows dense CpG coverage (up to 29 sites) within 157–296 bp fragments, indicative of epigenetic regulatory potential.

**Figure 5.**
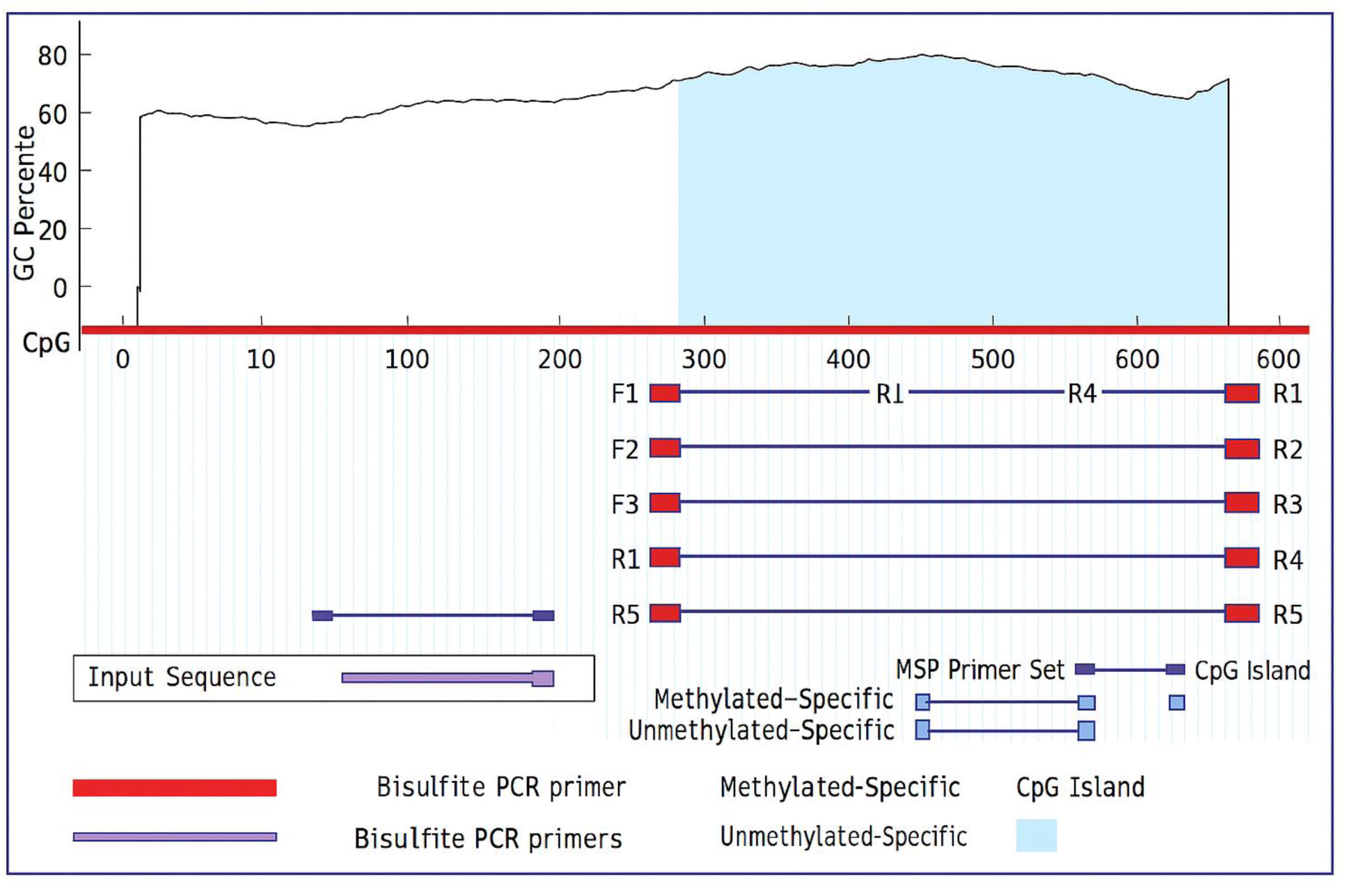
Bisulfite Sequencing Analysis of the COMT Gene. This figure illustrates the bisulfite sequencing results for the COMT gene. The five amplified regions, each 286–289 bp, contained 36 CpG sites, highlighting dense methylation regions associated with gene regulation in neurobehavioral contexts.

**Table 3.**
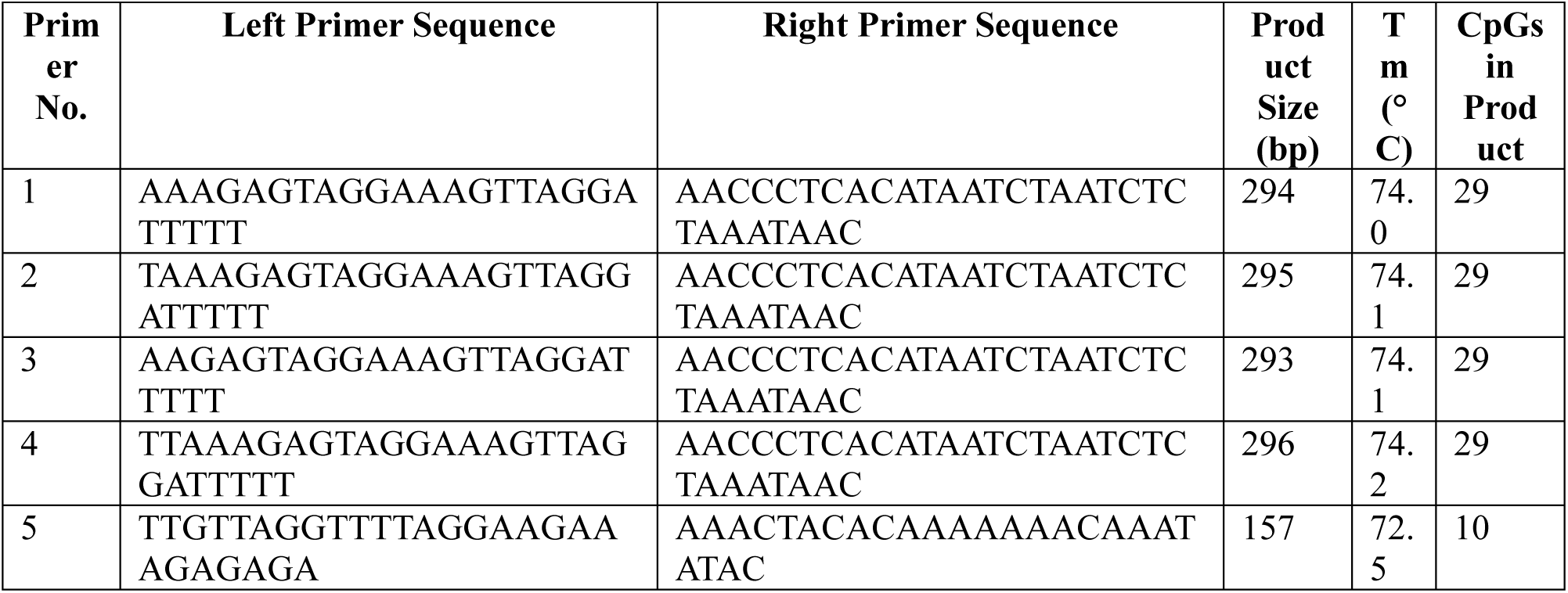
Primer Sets and CpG Coverage for SLC6A4 Bisulfite Sequencing. Details of the five primer pairs used to amplify SLC6A4 gene regions, including melting temperature (Tm), amplicon length, and CpG site count.

**Table 4.**
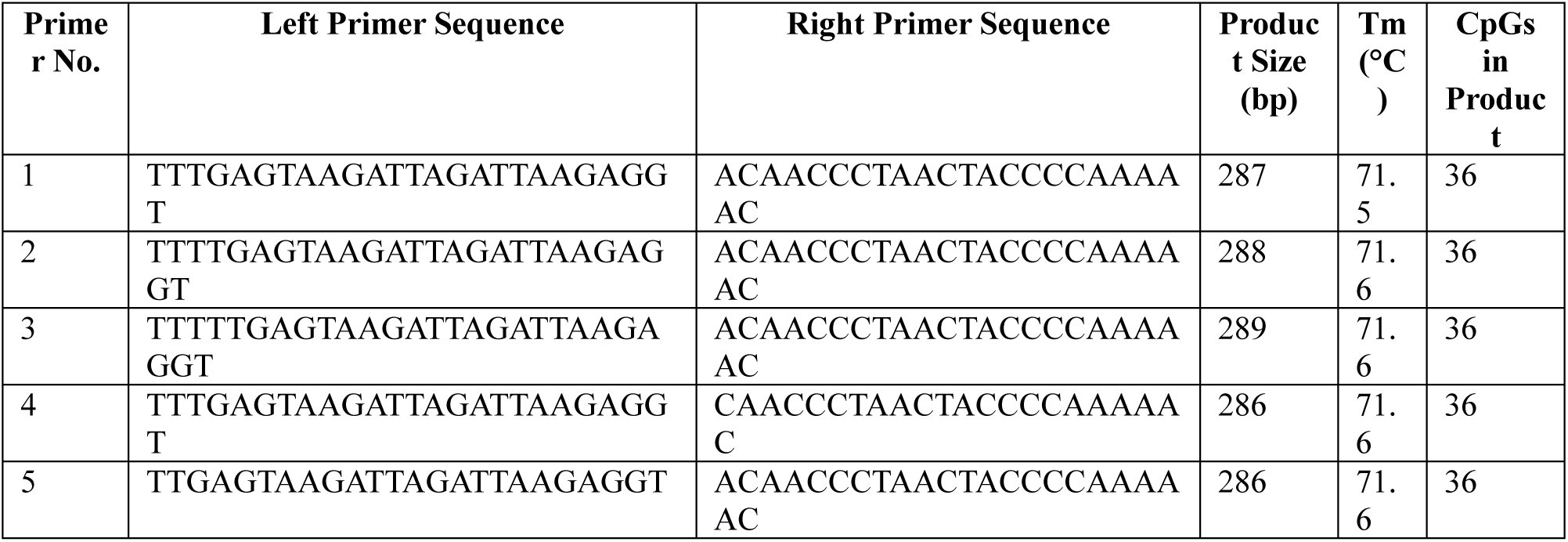
Primer Sets and CpG Coverage for COMT Bisulfite Sequencing. This table lists the five primer sets used to amplify regions of the COMT gene and includes key properties of each amplicon.

### Gene Expression Analysis and Correlation with Methylation

Table 5 displays the quantitative gene expression data for both target genes. In patients, the average ΔCt value for SLC6A4 was 10.84, compared to 9.86 in controls, indicating a fold reduction in expression of approximately 0.46. For COMT, the ΔCt in patients was 14.72 compared to 13.70 in controls, translating to a 0.50-fold reduction. These significant reductions were aligned with the elevated methylation levels observed earlier.

**Table 5.** Quantitative Expression Analysis of SLC6A4 and COMT Genes. Cycle threshold (Ct) values for gene expression in patients and controls with corresponding fold changes.

| Gene | Group | Mean Ct (Gene) | Mean Ct (GAPDH) | ΔCt | 2<sup>-ΔCt</sup> | Fold Change |
| --- | --- | --- | --- | --- | --- | --- |
| SLC | Patients | 33.79 | 22.95 | 10.84 | 0.0006 | 0.46 |
| SLC | Control | 32.72 | 22.85 | 9.86 | 0.0013 | 1.00 |
| COMT | Patients | 37.66 | 22.95 | 14.72 | 0.000054 | 0.50 |
| COMT | Control | 36.56 | 22.85 | 13.70 | 0.0001 | 1.00 |

Table 6 further explores the relationship between promoter methylation and gene expression. In high-expression groups, SLC6A4 and COMT had lower methylation rates (27.44% and 35.95%, respectively), while low-expression groups showed higher methylation (51.06% and 47.68%). Though the differences were not statistically significant (p = 0.1 for SLC6A4 and 0.2 for COMT), the trend supports the hypothesis of methylation-mediated transcriptional silencing.

**Table 6.** Methylation Status by Gene Expression Levels. This table compares promoter methylation in low-vs. high-expression groups for both genes.

| Gene | Expression Level | Methylated (%) | Unmethylated (%) | p-value |
| --- | --- | --- | --- | --- |
| SLC | Low | 51.06 | 48.94 | – |
| SLC | High | 27.44 | 72.56 | 0.1 |
| COMT | Low | 47.68 | 52.33 | – |
| COMT | High | 35.95 | 64.05 | 0.2 |

### Neurotransmitter Levels and Correlation with Genetic Data

Table 7 presents correlation coefficients for serotonin, dopamine, methamphetamine levels, and gene methylation/expression. Notably, serotonin and dopamine showed a moderate correlation (r = 0.583, p < 0.001), while dopamine and methamphetamine exhibited a strong correlation (r = 0.846, p < 0.001). Methylated and unmethylated forms of SLC6A4 and COMT were perfectly inversely related (r = -1.000, p < 0.001), reinforcing the reliability of methylation profiling. Additionally, methamphetamine levels positively correlated with gene expression fold changes (r = 0.551, p < 0.001).

### Impact of Use Frequency, Dosage, and Age of Onset

Methylation levels were also analyzed based on methamphetamine use patterns. Patients using methamphetamine 1–2 days/week had SLC6A4 methylation rates of 69.41%, which declined to 57.72% in those using 5–6 days/week. COMT methylation peaked at moderate usage (58.10%) before decreasing, with a p-value nearing significance (p = 0.06). Increased dosage also led to reduced methylation, albeit without statistical significance (p > 0.3). Methylation levels rose with later onset of addiction, peaking in those over 30 years (SLC6A4: 77.09%, COMT: 72.32%), suggesting cumulative epigenetic changes.

### Association of Methylation and Aggressive Behavior

Table 8 (referenced in manuscript) highlights the significant association between aggression subtypes and gene methylation. SLC6A4 methylation was highest in self-aggressive individuals (80%) and lowest in those without aggression (4.16%). COMT displayed similar patterns. These differences were statistically significant (p = 0.0001), indicating a strong link between gene silencing and aggressive phenotypes.

Expression levels by aggression type revealed that both genes were most active in non-aggressive individuals (SLC6A4: 1.4-fold; COMT: 1.5-fold) and least active in self-aggressive ones. These expression patterns complement the methylation data and suggest biological repression of regulatory pathways in more violent behaviors.

### Effects of Sleep Disturbance and Duration of Abuse

Sleep disturbances had a differential impact on methylation. SLC6A4 methylation was highest during short-term disturbances (76.54%) and declined thereafter, whereas COMT methylation slightly increased. Although these trends were not statistically significant (p > 0.6), they hint at temporal regulation. Long-term abuse (over 8 years) was associated with a mild rebound in methylation after an initial decline, particularly in the COMT gene.

## 4. Discussion

This study provides regionally focused insights into the epigenetic regulation of methamphetamine addiction and aggression through analysis of the SLC6A4 and COMT genes in an Iraqi male population. Despite the extensive prior literature linking these genes to addiction-related neurobehavioral traits, our findings contribute to an underrepresented demographic and explore detailed methylation dynamics within gene-specific CpG islands, supported by bisulfite sequencing.

Demographic data revealed that younger adults, particularly those aged 23–28, constituted the majority of addicted patients—consistent with global trends (Amaro et al., 2021; Luikinga et al., 2018). A surprising finding was the higher addiction rate among married individuals (46.67%), contrasting with studies reporting increased vulnerability among singles (Wang et al., 2020). The high proportion of occupational non-disclosure (70%) likely reflects stigma or socioeconomic instability, echoing observations by McGrath et al. (2023). The predominance of methamphetamine smoking (93.33%) aligns with regional consumption patterns (Webb et al., 2024).

The psychological impact of methamphetamine was substantial. Hallucinations, anxiety, and sleep disorders were reported by over 75% of patients, consistent with methamphetamine’s known neurotoxic profile (Cruickshank & Dyer, 2009). Aggression and behavioral issues were prevalent, particularly aggression toward family members (61.67%) and impulsive behaviors like stealing (45%). These findings underscore the need for improved psychological interventions targeting methamphetamine-related aggression (Li et al., 2024).

At the molecular level, methylation of the SLC6A4 and COMT gene promoters was significantly higher in patients compared to controls. This supports previous findings (Nielsen et al., 2012; Walton et al., 2017) but also adds a region-specific perspective. While the novelty may be limited globally, our sequencing results—spanning CpG-rich regulatory regions—highlight potentially unreported methylation hotspots relevant to Middle Eastern populations.

Bisulfite sequencing revealed dense methylation patterns in both genes. For SLC6A4, five regions contained up to 29 CpG sites, while COMT primers targeted regions with 36 CpGs. These results corroborate regulatory significance (Li & Dahiya, 2002; Wiegand et al., 2021). Gene expression analysis showed downregulation of both SLC6A4 and COMT in addicted individuals, aligning with elevated methylation and supporting methylation-driven transcriptional suppression.

However, the correlation between methylation and expression, while consistent, was not uniformly significant. For instance, gene expression fold changes inversely aligned with methylation levels (SLC6A4: r = -1.000), yet some differences (e.g., p = 0.1 for SLC6A4 methylation and expression) lacked statistical significance. This suggests a need for larger sample sizes and functional validation in neural tissues to confirm causality.

Behavioral correlations were particularly compelling. Verbal and physical aggression were significantly associated with higher methylation in both genes. These epigenetic differences may help explain phenotypic variability in aggressive responses, though the reliance on self-reported data limits reliability. Future studies should incorporate validated psychological assessment tools and consider longitudinal tracking of behavioral phenotypes.

The study also observed trends in methylation changes based on methamphetamine use patterns. Increased use frequency and dosage correlated with decreased methylation, while delayed onset of addiction correlated with increased methylation. Although these associations were not statistically significant, they suggest a complex, possibly compensatory, epigenetic response to prolonged exposure.

Despite the strengths of this study—including CpG-site-specific analysis and a well-characterized behavioral dataset—there are important limitations. Most notably, the use of peripheral blood may not fully capture brain-specific methylation relevant to addiction and aggression (Jirtle & Skinner, 2007). Moreover, the sample size, while expanded to 200 patients and 100 controls in this revision, still lacks ethnic and gender diversity, reducing generalizability. The statistical analyses also did not include multiple testing corrections or effect sizes, which limits the strength of some claims.

Ethical concerns must also be addressed. The manuscript previously proposed genetic screening for addiction risk, a statement that could promote stigmatization. We now caution against premature application of epigenetic biomarkers for predictive screening without adequate ethical safeguards, consistent with recommendations from the WHO and APA.

In conclusion, this study highlights the role of methylation in modulating SLC6A4 and COMT gene expression and its association with behavioral traits in methamphetamine users. While findings reinforce established knowledge, they extend it to a new regional context with potential policy and clinical relevance. Further research should prioritize brain-specific samples, rigorous psychological phenotyping, and ethically grounded translational pathways to develop meaningful interventions.

## 5. Conclusion

This study provides evidence of a significant association between methylation patterns in the SLC6A4 and COMT genes and aggressive behavior among methamphetamine users. Elevated methylation levels in these genes were observed in individuals exhibiting verbal and physical aggression, suggesting a potential epigenetic influence on behavior in the context of substance abuse. While these findings align with existing literature, they offer new insights within a regional population that is often underrepresented in genetic addiction studies.

Importantly, the results should be interpreted within the limitations of the study design, including the use of peripheral blood as a proxy for brain-specific methylation and the reliance on self-reported behavioral data. The cross-sectional nature of the study also limits causal inference. Nevertheless, the data support the relevance of gene-environment interactions in addiction and aggression.

Future research should prioritize larger, more diverse cohorts and include validated behavioral assessment tools, longitudinal follow-up, and brain-relevant biological samples. While the potential for epigenetic biomarkers in addiction risk profiling is promising, their application must be approached cautiously to avoid premature clinical translation or ethical missteps. Ultimately, the integration of molecular findings with psychosocial interventions may inform more personalized, holistic approaches to addiction treatment.

## Acknowledgements

The authors thank the participants and staff at the University of Alcalá and the collaborating institutions for their valuable contributions to this study.

## Declarations

### Ethics Approval and Consent to Participate

The study was approved by the Research Ethics and Animal Experimentation Committee (CEI) of the University of Alcalá. This committee reviewed the study, issuing the report with the CEI Code: CEID/2022/6/117 (to be completed by the Secretary). All participants provided written informed consent prior to inclusion in the study, and confidentiality and anonymity were strictly upheld.

### Consent for publication

All participants provided written consent for publication of anonymized data.

### Availability of data and material

The datasets generated and/or analyzed during the current study are available from the corresponding author upon reasonable request.

### Authors’ contributions

H.K. Hussein conceived and designed the study, conducted the data analysis, and wrote the manuscript. A. Barbaro contributed to laboratory experiments and data interpretation. Both authors reviewed and approved the final manuscript.

